# The Growth of *Xiphophorus hellerii* is Negatively Correlated with Colony Density

**DOI:** 10.1101/2025.09.22.677796

**Authors:** Lindsey Sanchez, Markita Savage, Will Boswell, Yuan Lu

**Affiliations:** Institute for Molecular Life Sciences, Texas State University

## Abstract

*Xiphophorus* fishes serve as important animal models in biomedical research. To investigate colony density influence on growth, we compared 6-, 9-, and 12-month-old fish raised in low- (1 fish/2L), mid- (1 fish/L), and high-density (2 fish/L) in re-circulating aquariums. As expected, the results indicate that fish reared in low-density aquariums exhibit the greatest growth, and fish reared in high-density aquariums result in the least growth.

## Introduction

*Xiphophorus* is a genus of 26 small freshwater fish species native to Mexico and Central America [1]. These fish are popular ornamental fish among hobbyists due to their vibrant and varied pigmentation patterns. In addition to their ornamental value, *Xiphophorus* species serve as animal models for studying evolution, ecology, development, behavior, genomics, toxicology, cancer, and metabolic diseases [2]. The strength of the *Xiphophorus* model is highlighted in biomedical research and has been recognized as an evolutionary medicine model [3-5].

*Xiphophorus* species exhibit morphological, behavioral, and genetic divergence [1, 2, 6]. Their ability to produce inter-species hybrids positions the *Xiphophorus* model in a unique niche for analyzing parental alleles co-segregated with morphological, behavioral, or molecular phenotypes. These gene-phenotype correlation analyses, referred to as an evolutionary genetics screen, represent a distinct forward genetic screening approach that has contributed to the understanding of multiple traits relevant to human diseases. Available *Xiphophorus* resources include live animals for most species, interspecies hybrids, reference genomes for the genus, reference transcriptome reflecting basal level, dynamic temporal changes for multiple organs, reference gut microbiome, reference single cell transcriptome and cell atlas, pathological images, preserved tissue samples, preserved germplasms, and publicly available genomic and transcriptomic data [2, 6-11].

These resources are hosted in the *Xiphophorus* Genetic Stock Center (XGSC) at Texas State University. Myron Gordon initially founded the XGSC at the American Museum of Natural History in New York City in 1926. Throughout the century, the fish stocks continued to expand and have been maintained using the same husbandry protocol. Briefly, mature males and females are kept in isolated static aquariums at a density of 1 fish/gallon of water until they are recruited for an experiment or placed in mating pairs. However, isolated static aquariums can differ in water chemistry between individual tanks, which can influence overall animal wellbeing, thus resulting in untraceable variations among animals. To minimize cross-aquarium variations, modern automated re-circulation systems have been implemented to allow all fish living in one system to be exposed to the same water chemistry. The usage of such a system enables investigations into influences of environmental cues on animal health.

In this study, we seek to understand the influence of colony density on *Xiphophorus* growth in a recirculating rack. Three groups, low, mid, and high-density groups, were created by placing the same number of 1-day-old *Xiphophorus* larvae into 18.9L, 37.8L, and 75.6L re-circulating aquariums. Body length and depth were measured at 6-, 9-, and 12-month for each density group. We found that colony density is negatively correlated with *Xiphophorus* growth.

## Results

A total of 107 swordtails were randomly assigned to high-density (n=34; 2 fish/1L), mid-density (n=36; 1 fish/1L), and low-density (n=37; 1 fish/2L) test groups. Throughout the 12-month study, one animal in the low-density group died. No disease or malformation was observed.

Per time point, lower density groups (i.e., mid-density vs. high-density; and low-density vs mid-density) exhibited statistically significant increase in body length and depth (Welch Two Sample t-test p<0.05), except 6-month mid-density vs. low-density comparison (Fig. 1). The ratios of body length over depth were consistent among all density groups (Fig. 1).

**Figure 1.**
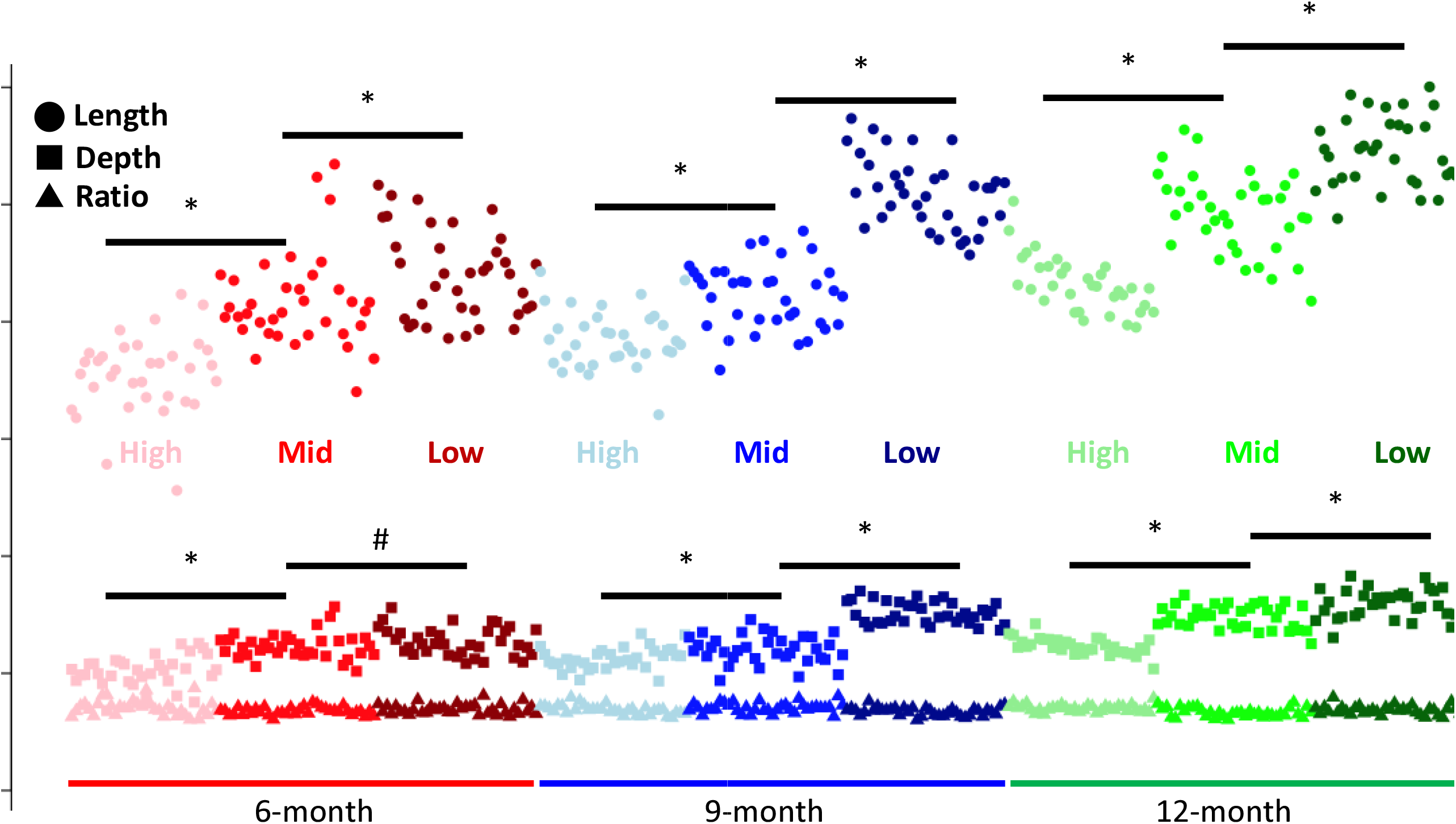
Standard length and depth measurements of fish reared in low-, mid-, and high-density aquariums. Measurement of fish standard length and depth (mm) is plotted for all density groups in all data points. * means p-value < 0.05, and # means p-value > 0.05.

Per density group, other than the mid-density group between 6- and 9-months that do not show either length or depth differences, all other groups show statistically significant 9-month vs. 6-month and 12-month vs. 9-month growth in length and depth (Fig. 2).

**Figure 2.**
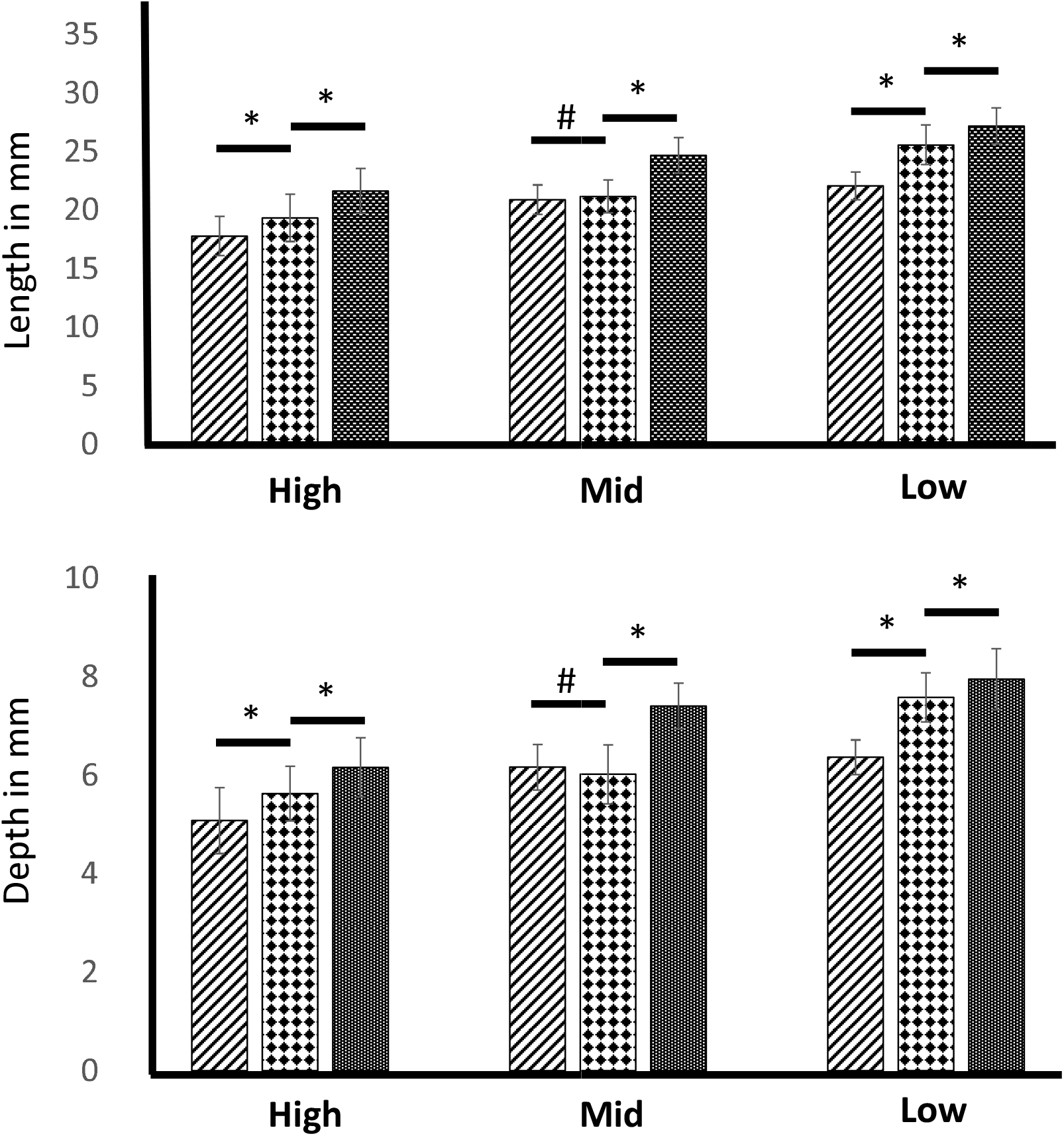
Standard length and depth comparison of fish reared in the same density group throughout time. Clustered bar graph shows mean values of standard length (top) and depth (bottom). Error bars represent standard deviation. Per bar cluster, mean values of 6-, 9-, and 12-month are shown from left to right. * means p-value < 0.05, and # means p-value > 0.05.

## Discussion

This study aimed to investigate the influence of *Xiphophorus* fish colony density on growth. Overall, the results clearly showed that high colony density is negatively correlated with body size. Ecological studies have shown that intra-population competition (i.e., higher colony density increases competition among members for limited resources, such as food) and increased foraging costs (higher colony density can deplete food sources and force individuals to forage over great distances, thereby increasing energy expenditure) can influence individual growth [12, 13]. In this study, different density groups were fed with equal amounts of food so that feeding competition could not explain the overall trend observed. Controversially, the only group of fish that may need to travel further for food are those in low-density groups, which showed the largest body size.

One plausible explanation is that chronic overcrowding causes increased stress levels that negatively affect individual growth and size. Similar findings, as shown in this study, have been observed in bumblebees [14], rats [15], rainbow trout [16], and *Simocephalus vetulus* [17]. Crowding can activate the hypothalamic-pituitary-adrenal (HPA) axis and lead to stress hormones such as cortisol [18]. Chronic crowding was reported to impact the weights of the thymus, liver, and endocrine glands [15], cause major epigenetic and gene expression changes [16], and induce chronic stress that suppresses growth hormone secretion [19-21].

In conclusion, we found that *Xiphophorus* fish exhibit a growth retardation related to non-nutritional colony density, likely due to potential chronic stress.

## Materials and Methods

### Animal Model

*Xiphophorus hellerii* used in this study are supplied by the *Xiphophorus* Genetic Stock Center. All fish were kept and samples taken in accordance with protocols approved by Texas State University Institutional Animal Care and Use Committee (IACUC 9048). A total of 107 one-day-old fry were produced by colony breeding of parentals and were randomly assigned to low-density (n=34), mid-density (n=36), and high-density (n=37). Fish in different density group were fed with the same amount of food that are adjusted to colony size, including Aquatox flake food and live brine shrimp twice a day.

By 6 months old, all fish were anesthetized with 0.01% MS-222. Upon loss of gill movement, standard length and depth measurements were taken using a digital caliper. Fish were then returned to their respective tanks. This process was repeated for 9 and 12-months-old.

### Statistical analyses

Data analyses were conducted using R (v4.3.1). Length and depth comparison between ages or density groups was conducted using the Welch Two-Sample t-test. P-value < 0.05 is used to determine statistical significance.

## Acknowledgements

This work was supported by the National Institutes of Health, Office of Director R24 OD-031467.

